# Phenotypic defects from the expression of wild-type and pathogenic TATA-Binding Proteins in new *Drosophila* models of Spinocerebellar Ataxia Type 17

**DOI:** 10.1101/2023.05.22.541820

**Authors:** Nikhil Patel, Nadir Alam, Kozeta Libohova, Ryan Dulay, Sokol V. Todi, Alyson Sujkowski

## Abstract

Spinocerebellar Ataxia Type 17 (SCA17) is the most recently identified member of the polyglutamine (polyQ) family of disorders, resulting from abnormal CAG/CAA expansion of TATA box binding protein (TBP), an initiation factor essential for of all eukaryotic transcription. A largely autosomal dominant inherited disease, SCA17 is unique in both its heterogeneous clinical presentation and low incidence of genetic anticipation, the phenomenon in which subsequent generations inherit longer polyQ expansions that yield earlier and more severe symptom onset. Like other polyQ disease family members, SCA17 patients experience progressive ataxia and dementia, and treatments are limited to preventing symptoms and increasing quality of life. Here, we report two new *Drosophila* models that express human TBP with polyQ repeats in either wild-type or SCA17 patient range. We find that TBP expression has age- and tissue-specific effects on neurodegeneration, with polyQ expanded SCA17 protein expression generally having more severe effects. In addition, SCA17 model flies accumulate more aggregation prone TBP, with a greater proportion localizing to the nucleus. These new lines provide a new resource for the biochemical characterization of SCA17 pathology and the future identification of therapeutic targets.

## INTRODUCTION

The polyglutamine (polyQ) family of progressive neurodegenerative disorders comprises a group of nine diseases caused by abnormal expansion of the trinucleotide repeat (CAG/A) that encodes glutamine (Zoghbi and Orr 2000; La Spada and Taylor 2003; Todi *et al*. 2007; Paulson *et al*. 2017; Lieberman *et al*. 2018; Buijsen *et al*. 2019; Liu *et al*. 2019). Spinocerebellar Ataxia Type 17 (SCA17) is the most recently discovered member of the polyQ disease family, resulting from CAG/CAA expansion in the gene that encodes the TATA box-binding protein (TBP), an essential player in the basal transcriptional machinery of all eukaryotes (Koide *et al*. 1999; Nakamura *et al*. 2001). SCA17 is largely an autosomal dominant inherited disorder (Fujigasaki *et al*. 2001), but *de novo* TBP mutations have also been reported (Bech *et al*. 2010) in patients.

Like other diseases in the polyQ family, SCA17 usually manifests in adulthood with classical symptoms that include ataxia, dystonia, and dementia (Toyoshima and Takahashi 2018; Liu *et al*. 2019). Brain imaging is characterized by cerebellar and brainstem atrophy and degeneration (Nakamura *et al*. 2001). SCA17 is unique, however, due to the extensive variability in both clinical presentation and age of onset, and the unusually small gap between normal and pathological polyQ repeat length (Toyoshima and Takahashi 2018; Gardiner *et al*. 2019; Liu *et al*. 2019). Wild-type alleles of TBP have 25-42 CAG/A repeats, while most SCA17 mutations fall within the repeat range of 46-55 (Gao *et al*. 2008). There is a weak correlation between repeat length and age of onset, with juvenile forms usually associated with polyQ lengths over 62, but several SCA17 studies find high genetic heterogeneity, even among the same family of patients (Liu *et al*. 2019). In fact, full disease presentation has been reported in patients with CAG/CAA repeat lengths as low as 41 (Origone *et al*. 2018).

Abnormal polyQ expansion in disease genes causes protein misfolding and aggregation, forming toxic intranuclear inclusions that lead to neuronal degeneration and death (Zoghbi and Orr 2000; Zoghbi and Orr 2009). In SCA17 patients, polyQ-expanded TBP accumulates in Purkinje neurons in the cerebellum, a finding that is recapitulated in animal models (Friedman *et al*. 2007; Ren *et al*. 2011). Studies in animal models have also discovered that polyQ expansion in TBP reduces endogenous protein levels and disrupts it normal function, yielding transcriptional dysregulation that further contributes to SCA17 disease progression (Friedman *et al*. 2008). Earlier *Drosophila* models employing hyperexpanded and amorphic protein expression have provided important information about TBP function, dysfunction, and interactions during SCA17 disease progression (Ren *et al*. 2011; Hsu *et al*. 2014), but until now there were no *Drosophila* models expressing human TBP with polyQ expansions within patient range.

Here, we report new *Drosophila* transgenic lines of SCA17 that express HA-tagged, full-length human TBP with CAG/CAA repeats encoding either 25Q (wild-type) or 63Q (SCA17). We find that global expression of either construct causes developmental arrest or delay, and that adult-restricted, ectopic expression of either Q25 or 63Q reduces survival and mobility whether the expression pattern is ubiquitous, in neurons, or in glia. In general, polyQ-expanded TBP expression is more toxic in adult flies and is associated with increased aggregation propensity of disease protein, particularly in mid-to late-life, compared to its wild-type version. This model adds to the toolkit of genetically isogenous *Drosophila* polyQ models, making possible rapid, cost-effective investigations and increasing our mechanistic understanding of SCA17 pathology.

## RESULTS

### Generation of wild-type and polyQ-expanded SCA17 models

We have generated several Gal4-UAS model lines that express human SCA disease proteins (Tsou *et al*. 2015; Tsou *et al*. 2016; Johnson *et al*. 2019), leveraging a cloning strategy that yields a single-copy, phiC31-dependent insertion into the same attp2 integration site, in the same orientation, on the third *Drosophila* chromosome (Groth *et al*. 2004). This expression system allows us to easily modify protein expression and makes it possible to compare across our *Drosophila* SCA models under conditions of similar protein expression levels and identical genetic background (Markstein *et al*. 2008; Ni *et al*. 2008; Tsou *et al*. 2016). As previously mentioned, wild-type TBP alleles range from 25 to 42 polyQ repeats, whereas SCA17 expansions typically extend from 46 to over 60 repeats, with CAA interruptions that may further stabilize disease penetrance (Liu *et al*. 2019). We used this strategy to generate two new *Drosophila* lines that express HA-tagged human TBP harboring repeat lengths of either Q25 (wild-type) or Q63 (SCA17), CAG/CAA polyQ tracts specifically designed to be within patient range (Toyoshima and Takahashi 2018) (Figure 1A). Genomic DNA was isolated from founder lines and the inserted sequences were amplified and sequenced, confirming integration into the third chromosome, in the correct insertion site and in the proper orientation (Figure 1B).

**Figure 1:**
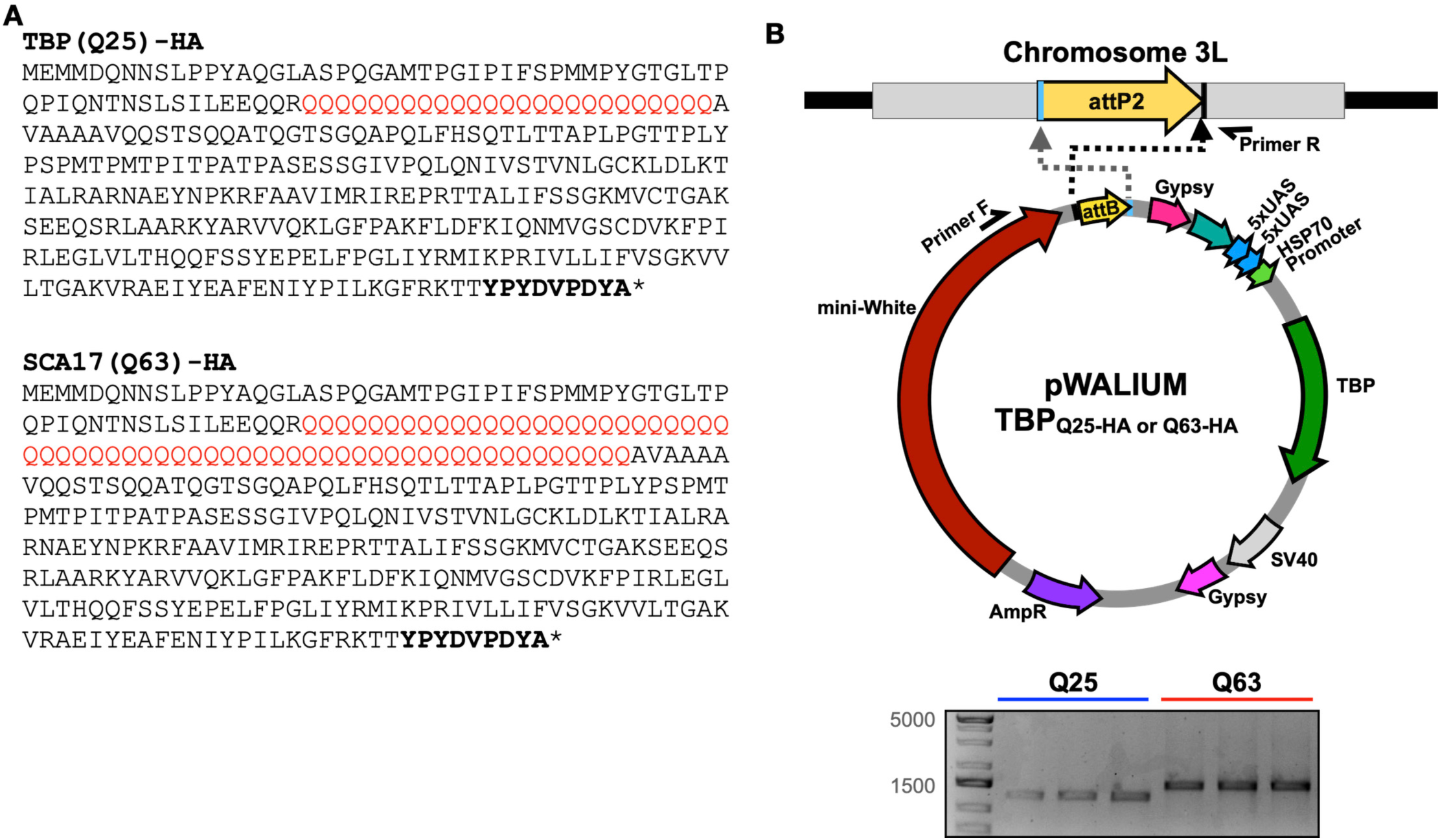
Generation of *Drosophila* SCA17 models. **(A)** Human TBP amino acid sequences used to model wild-type (Q25, upper), and pathogenic (Q63, lower) TBP, with polyQ mutations indicated in red. C-terminal HA tag in bold. **(B)** Top: Schematic representation of the cloning strategy to insert TBP cDNA into pWALIUM10.moe. Bottom: Triplicate PCR from genomic DNA indicating that both transgenes were integrated into the correct insertion site in the proper orientation.

### Developmental toxicity in TBP-expressing flies

TBP is a widely expressed protein. Thus, we first examined the toxicity of each transgene using the constitutive, ubiquitous driver sqh-Gal4. Unlike the majority of SCAs that typically manifest in adulthood, the age of onset for SCA17 patients ranges from 3 to 60 years (Toyoshima and Takahashi 2018). Constitutive, ubiquitous expression of either Q25 or Q63 in *Drosophila* led to marked developmental lethality compared to control flies (Figure 2A). Nearly all Q25 flies died at ‘pharate’ pupal stage, without emerging as adults, while ubiquitous Q63 expression caused high percentages of both embryonic and pupal lethality, with fewer than 10% of flies surviving to adulthood (Figure 2A). Interestingly, more Q63 flies survived to adulthood than those expressing Q25, suggesting a dominant negative effect of exogenous, wild-type TBP expression that may interfere with basal transcriptional machinery critical for development.

**Figure 2:**
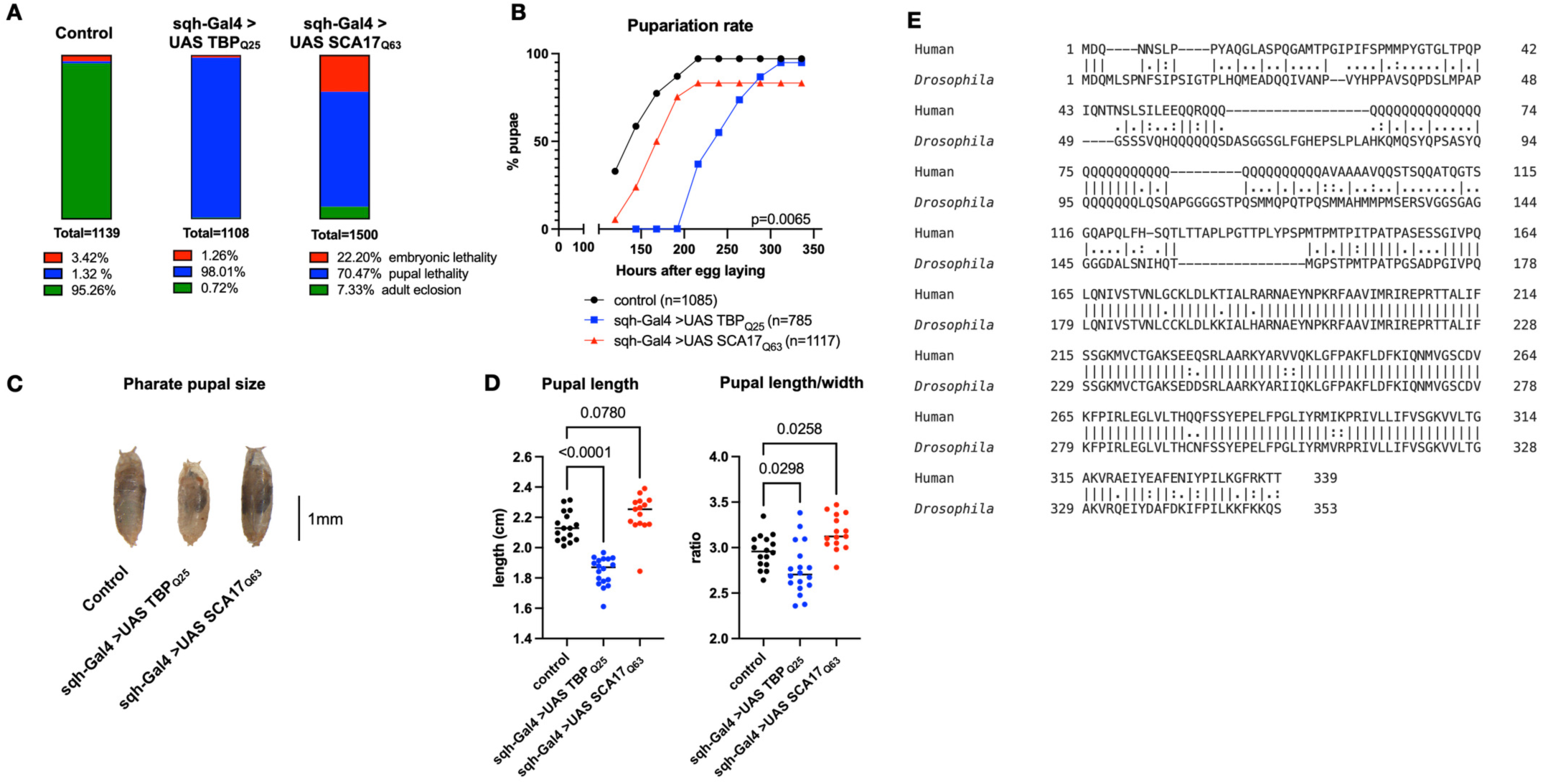
Developmental impact of ubiquitous TBP expression. **(A)** Percentage embryonic (red) and pupal (blue) lethality, and adult eclosion (green). Data from 3 individual crosses, n values indicated in panel. **(B)** Pupariation rate for control (black) wild-type (blue) and SCA17 (red) model flies. Data from 3 individual crosses. Statistics: linear regression, n values indicated in panel. **(C)** Representative image of pharate pupae, quantified in **(D)**. n≥15, Statistics: ANOVA with Dunnett’s post-hoc comparison. **(E)** Human-*Drosophila* TBP amino acid sequence alignment. **\*** (asterisk) indicates positions which have a single, fully conserved residue, : (colon) indicates conservation between groups of strongly similar properties

Development was delayed in both models compared to background control flies, with Q63 expression extending time to pupariation by approximately 1 day and Q25 expression nearly doubling pupariation time compared to controls (Figure 2B). Perturbations in maturation timing are often associated with disruptions in organismal growth (Delanoue and Romero 2020), and we observed small pupal size in Q25 flies (Figure 2C, D), while pupal length-to-width ratio was greater than controls in flies ubiquitously expressing Q63 (Figure 2C, D). Previous *Drosophila* lines with exogenous human TBP expression showed disrupted intrinsic TBP transcriptional function (Hsu *et al*. 2014), supporting a link between dysfunctional TBP and SCA17 pathogenesis. This finding is partially supported by the high structural and functional similarity of TBP in both flies and humans (Figure 2E). Altogether, these results highlight the critical importance of TBP sequence integrity and expression level to normal development, with perturbations in exogenous wild-type TBP expression severely impacting developmental timing and growth in these fly models.

### Constitutive TBP expression reduces adult lifespan and mobility, and impacts fly eye uniformity

We next assessed longitudinal physiology in our Q25 and Q63 flies that successfully emerged as adults. Expression of human polyQ proteins in *Drosophila* neurodegeneration models tends to reduce survivorship and impair mobility, phenocopying disease presentation in human patients (Blount *et al*. 2014; Tsou *et al*. 2015; Tsou *et al*. 2016; Sutton *et al*. 2017; Ristic *et al*. 2018; Johnson *et al*. 2019; Walters *et al*. 2019; Johnson *et al*. 2022; Prifti *et al*. 2022; Sujkowski *et al*. 2022). Compared to age-matched background controls, ubiquitous Q25 expression significantly decreased survival in both female and male flies, and survival in Q63 flies was reduced even further (Figure 3A). Similarly, mobility was severely impaired across ages in both female (Figure 3B) and male (Figure 3C) Q25 flies, while Q63 expression impacted mobility more severely, with most flies not responding to the climbing stimulus and remaining on the bottom of the vial (Figure 3D).

**Figure 3:**
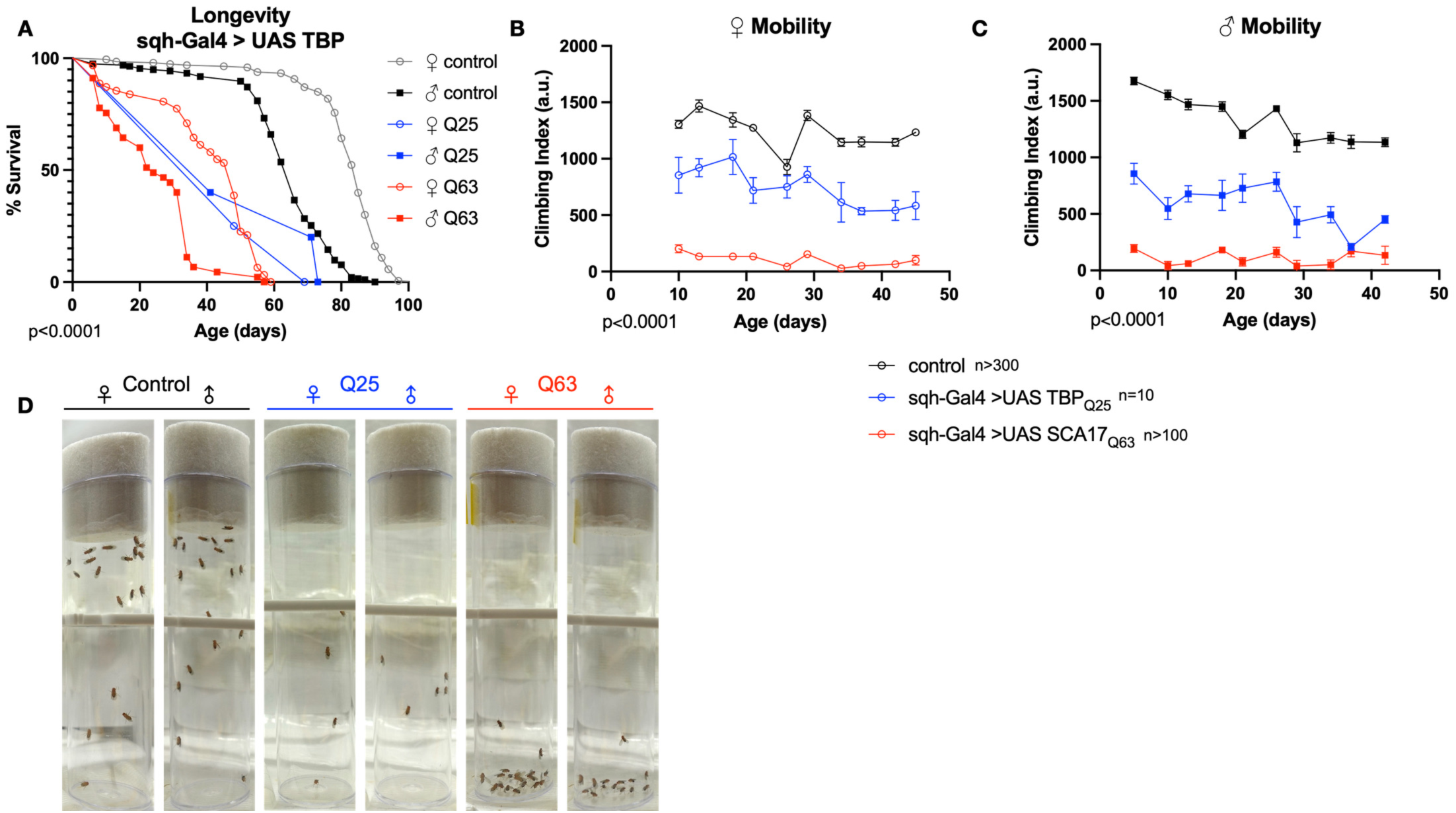
Effects of ubiquitous TBP expression on *Drosophila* lifespan and mobility. Impact of constitutive, ubiquitous TBP Q63 (red) or TBP Q25 (blue) expression on survival **(A)** and climbing speed in female **(B)** and **(C)** male flies compared to age-matched controls (black). Open symbols, females, closed symbols, males. Statistics: Survival, log-rank, excluding sqh- Gal4 > UAS TBP_Q25_ (low n, see methods). Motility: Linear regression for slope, intercept. The same flies were used for both mobility and survival. n values indicated on panels.

The *Drosophila* eye has been used extensively to assess neural toxicity in a variety of neurodegenerative disease models (St Johnston 2002; Matsumoto *et al*. 2003; Passarella and Goedert 2018; Mishra and Knust 2019; Smylla *et al*. 2021). We therefore used GMR-Gal4 to drive expression of each construct specifically in eyes, observing for phenotypic neurodegeneration with a scoring system we have used before (Figure 4A). Neither construct caused detectable anomalies in the external eye in one week old flies (Figure 4B, upper panels), but progressive defects were observed in both Q25 and Q63 expressing flies by week 5 (middle panels) that worsened by 10 weeks of age and were more severe in fly eyes expressing Q63 (Figure 4B bottom panels, quantifications in Figure 4C). Taken together, these results indicate that both wild-type (Q25) and polyQ expanded (Q63) human TBP expression is toxic in male and female *Drosophila*, with Q25 expression having pronounced effects on normal development and Q63 expression more severely impacting longevity, motility, and eye toxicity.

**Figure 4:**
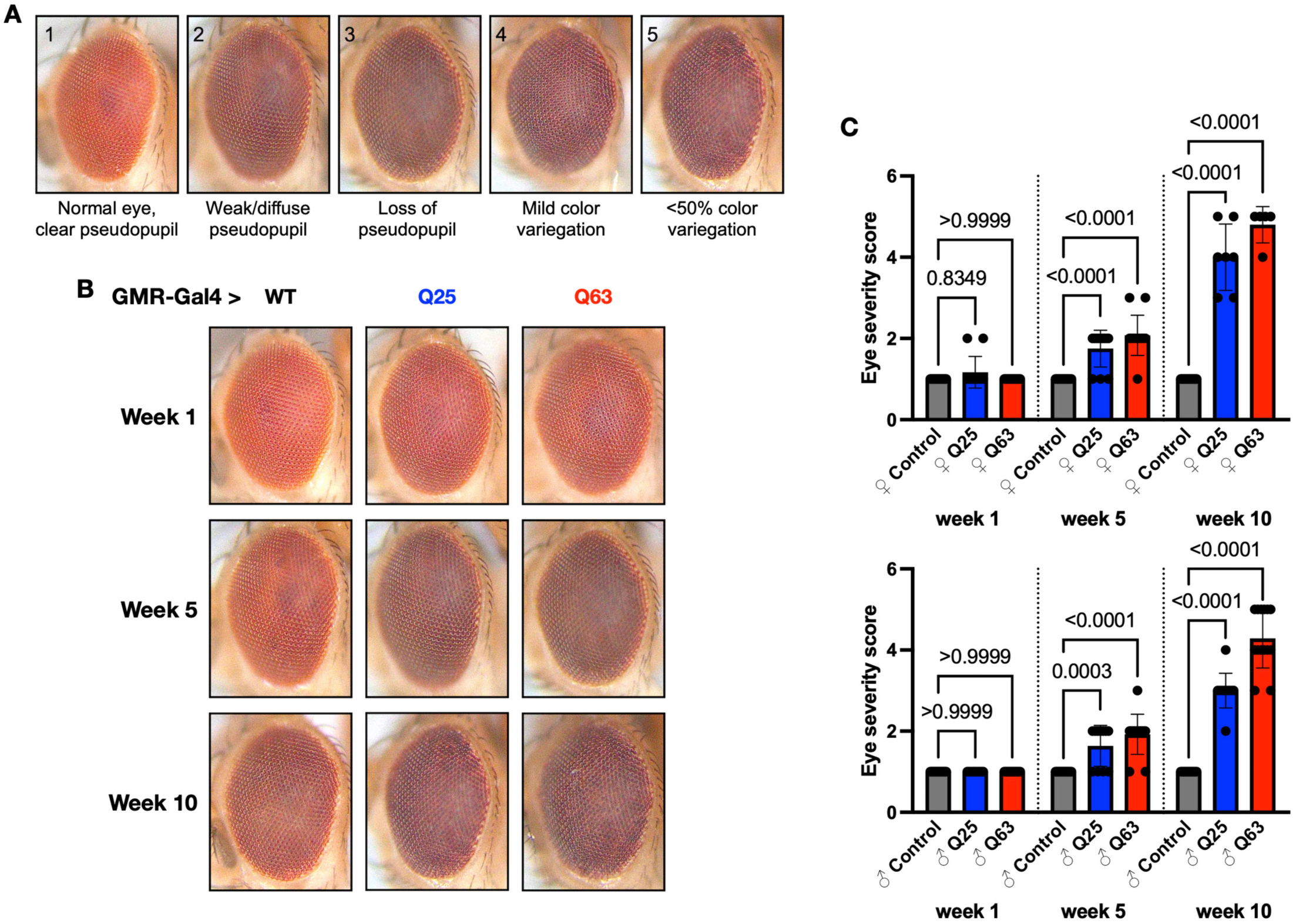
Effects of TBP expression in *Drosophila* eyes. **(A)** Scoring parameters. **(B)** Representative images of adult eyes in control flies (black) and flies expressing either Q25 (blue), or Q63 (red) in eyes at the indicated ages, quantified in **(C).** Statistics: ANOVA with Dunnett’s post-hoc comparison. Bars indicate mean -/+ SD, n≥10.

### Adult-specific TBP expression in Drosophila

In order to circumvent the high developmental lethality of our constructs, we next used the inducible Gal4-UAS Gene-Switch system (Roman and Davis 2002) to restrict TBP expression to adult tissues. We and others have shown that polyQ proteins increase in amount and aggregation propensity with age, and that disease protein aggregation is associated with toxicity (Blount *et al*. 2014; Tsou *et al*. 2015; Tsou *et al*. 2016; Sutton *et al*. 2017; Ristic *et al*. 2018; Johnson *et al*. 2019; Blount *et al*. 2020; Johnson *et al*. 2022; Sujkowski *et al*. 2022). Flies expressing either Q25 or Q63 in all adult tissues (RU+ (ON), Figure 5A, D, Figure S1A, D), had strong TBP expression in weeks 1, 3, and 5 of adult life. To assess aggregation propensity, we resolved both Q25 and Q63 proteins through SDS-PAGE and quantified SDS-soluble versus SDS resistant species across timepoints. In both female (Figure 5A, D) and male (Figure S1A, D) Q63 flies, SDS-resistant protein increased in weeks 3 and 5. In contrast, Q25 expression did not significantly alter SDS-resistant protein aggregation at any age.

**Figure 5:**
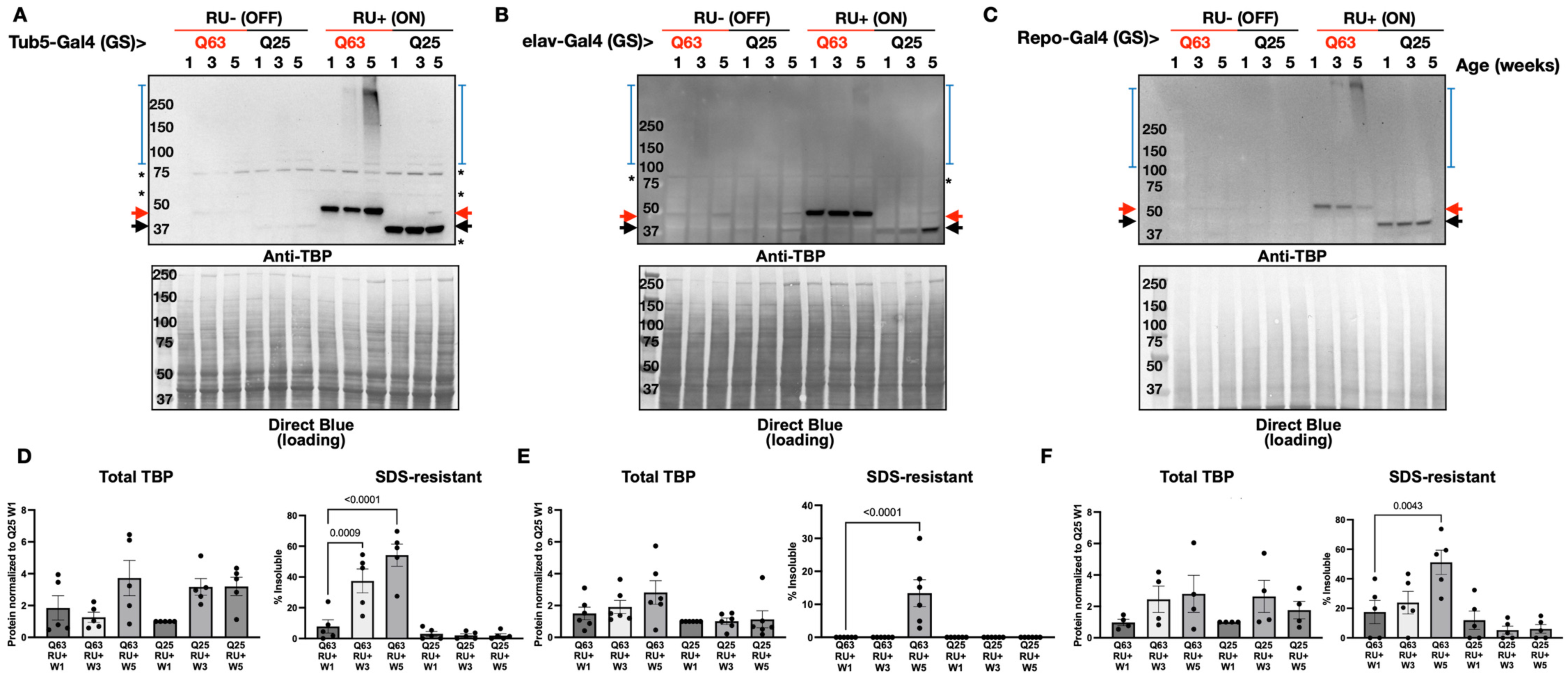
Impact of aging on TBP aggregation. Representative Western blots in uninduced female control (RU-(OFF)) and experimental flies (RU+ (ON)) with adult-specific, pathogenic TBP(Q63, red) or wild-type (Q25, black) TBP expression in weeks 1, 3, and 5 of adulthood, indicated by lane. Expression was induced on adult day 3 in **(A)** all tissues **(quantified in D)**, neurons **(B, E)**, or glia **(C, F)** and continued until sample isolation. Black arrows: TBP, Red arrows: polyQ expanded TBP, blue brackets: SDS-resistant TBP, asterisks: non-specific signal. Quantification and statistics: ANOVA with Dunnett’s post-hoc comparison. Bars indicate mean - /+ SD, n=5 biological replicates of 3 flies per lysate.

We observed similar patterns of TBP accumulation and aggregation when expression was restricted to either adult neurons (Figure 5B, E, S1B, E) or adult glia (Figure 5C, F, S1C, F). In both female and male flies, Q25 and Q63 expression was strongly detected across ages, and aggregation propensity increased in Q63 flies whether protein expression was restricted to adult neurons or glia, with the greatest increase in SDS-resistant protein accumulation at week 5. Our results indicate that although overall protein levels of either construct do not necessarily increase with age, Q63 protein is more prone to progressive aggregation whether expressed in adult neurons, adult glia, or in all adult tissues.

### Tissue specific adult TBP expression negatively impacts aging physiology

We next assessed tissue- and time- dependent effects of TBP expression on longevity and mobility using the same inducible drivers as above. First, we examined the effects of global, adult-specific expression using Tub5-Gal4 (GS), since TBP is ubiquitously expressed throughout the body (Liu *et al*. 2019). Adult specific expression of either Q25 (Figure 6A, B) or Q63 (Figure 6C, D) reduced lifespan in female flies without significant effects on mobility.

**Figure 6:**
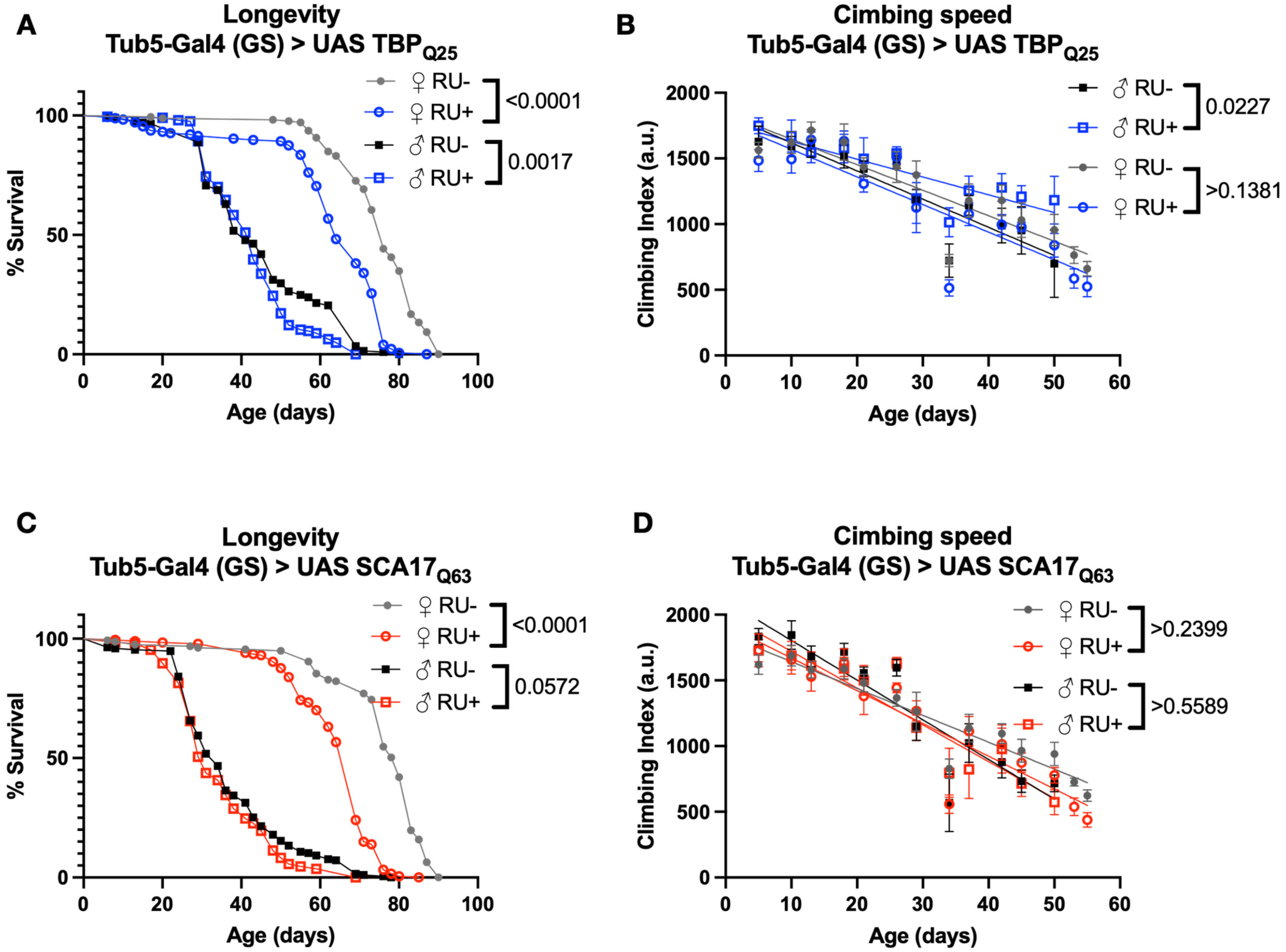
Impact of adult specific, ubiquitous TBP expression on longevity and motility. Representative survival **(A, C)** and climbing speed **(B, D)** experiments in male and female flies expressing Q25 **(blue, A, B)** and Q63 **(red, C, D)** TBP in all adult tissues compared to age-matched, uninduced control siblings (closed symbols). The same flies were used for both survival and climbing assays and assessed longitudinally. Statistics: Survival: log-rank, climbing speed: linear regression. Longevity and motility experiments performed in 2 biological replicates, n≥186.

Like other members of the polyQ disease family, neurons are particularly susceptible to degeneration and cell death despite TBP being widely expressed (Liu *et al*. 2019). Adult specific, pan-neural Q25 expression using elav-Gal4 (GS) did not affect either lifespan (Figure 7A) or mobility (Figure 7B) in comparison to uninduced background control flies. To further explore the connection between TBP expression and SCA17 disease phenotypes we then examined intracellular TBP localization. One common pathological hallmark of polyQ expansion diseases is the formation of neuronal intranuclear inclusions (Lieberman *et al*. 2019), a finding recapitulated in both cell and animal models of SCA17 (Friedman *et al*. 2007; Huang *et al*. 2011). In flies expressing Q25 specifically in adult neurons, histological examination of the adult neuromuscular junction (NMJ) showed TBP localization in both the nucleus and cytoplasm of muscle cells, with approximately half of the nuclei positive for TBP (Figure 7E, F).

**Figure 7:**
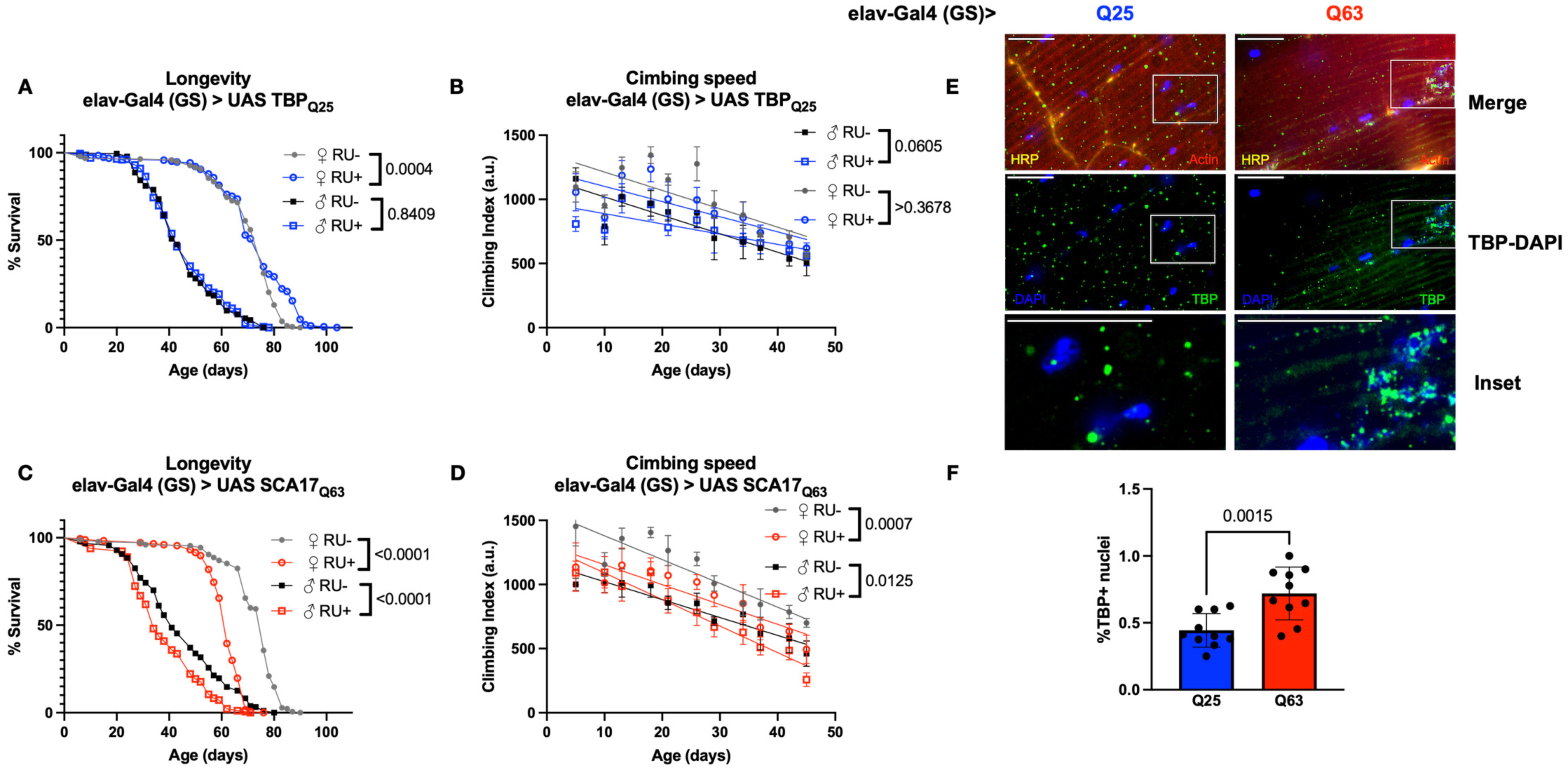
Effect of adult specific neural TBP expression on neuromuscular physiology. Representative survival **(A, C)** climbing speed **(B, D)** and experiments in male and female flies expressing Q25 **(blue, A, B)** and Q63 **(red, C, D)** TBP in adult neurons compared to age-matched, uninduced control siblings (closed symbols). The same flies were used for both survival and climbing assays and assessed longitudinally. Representative 100X images of adult NMJ for Q25 **(E)** and Q63 flies **(F)**. Red: actin, Yellow: HRP, Green: TBP, Blue: DAPI. Scale bar=15μm. Statistics: Survival: log-rank, climbing speed: linear regression. Longevity and motility experiments performed in 2 biological replicates, n≥184. Histology, n=10.

Unlike TBP Q25, expression of TBP Q63 specifically in adult neurons shortened lifespan in both female and male flies (Figure 7C), and significantly reduced mobility compared to age-matched, uninduced controls (Figure 7D). In addition, our neural Q63 model had a greater proportion of intranuclear TBP colocalization than Q25 flies (Figure 7E,F).

We then repeated these studies using Repo-Gal4 (GS) to restrict TBP expression to adult glia. We and others have demonstrated that polyQ expansion in glial cells contributes to progressive neurodegenerative phenotypes in animal models (Furrer *et al*. 2011; Yang *et al*. 2017; Johnson *et al*. 2022; Schuster *et al*. 2022) and may also participate in human SCA17 pathogenesis (Yang *et al*. 2017). Both Q25 and Q63 expression in adult glia decreased longevity (Figure 8A, C) and impaired mobility (Figure 8B, D), with glial-specific Q63 model females affected most severely (Figure 8C, D). Flies expressing either Q25 or Q63 in adult glia accumulated TBP intranuclearly, with the highest proportion of colocalization observed in our glial models of Q63 (Figure 8E, F).

**Figure 8:**
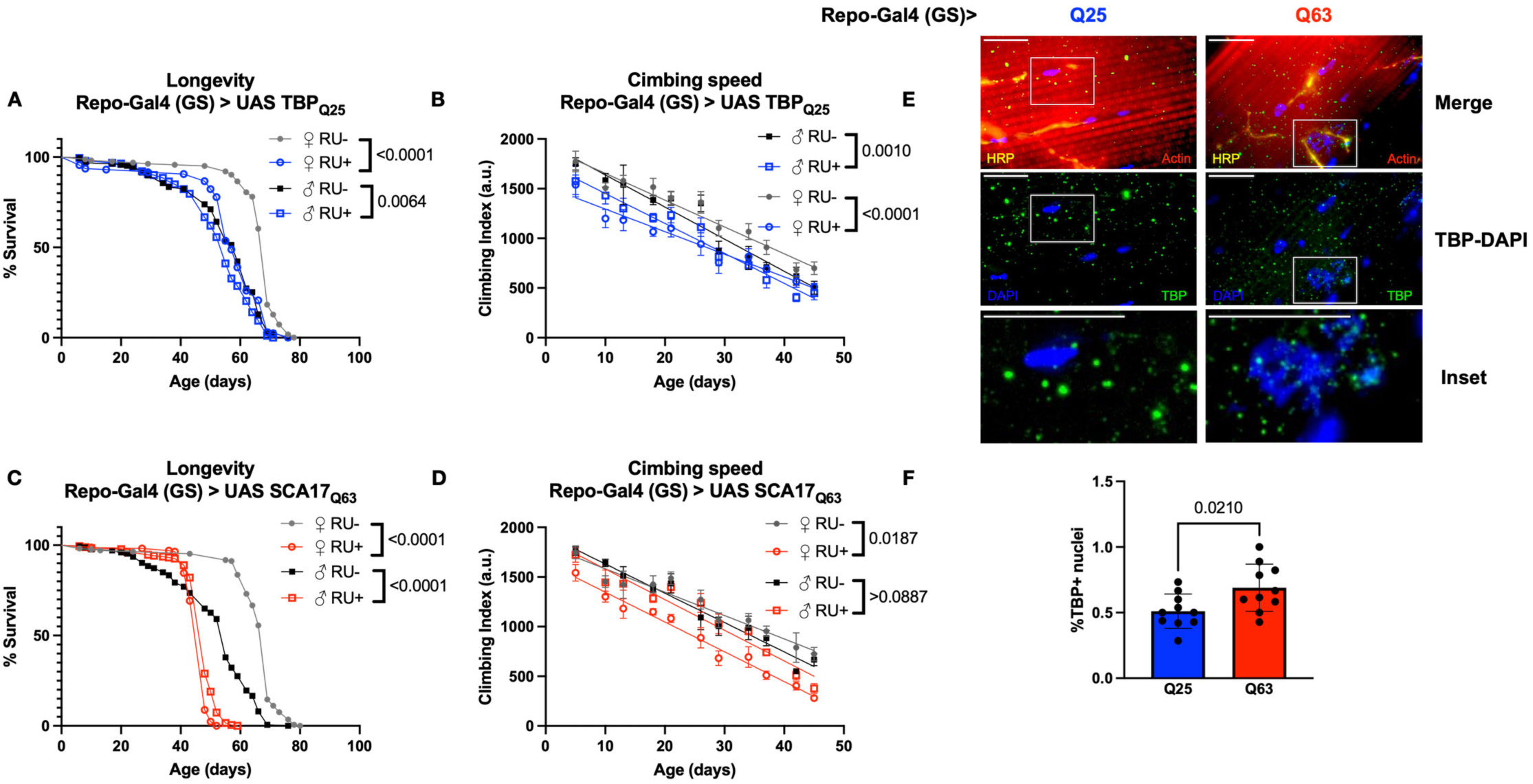
Effect of adult specific glial TBP expression on neuromuscular physiology. Representative survival **(A, C)** climbing speed **(B, D)** and experiments in male and female flies expressing Q25 **(blue, A, B)** and Q63 **(red, C, D)** TBP in adult glia compared to age-matched, uninduced control siblings (closed symbols). The same flies were used for both survival and climbing assays and assessed longitudinally. Representative 100X images of adult NMJ for Q25 **(E)** and Q63 flies **(F)**. Red: actin, Yellow: HRP, Green: TBP, Blue: DAPI. Scale bar=15μm. Statistics: Survival: log-rank, climbing speed: linear regression. Longevity and motility experiments performed in 2 biological replicates, n≥173. Histology, n=10.

These new *Drosophila* lines allow for future biochemical characterization of the relative contribution of tissue-specific polyQ expansion to SCA17 disease progression. In SCA17, like other polyQ family members, certain neural populations are more sensitive to dysfunction and death despite disease proteins being widely expressed. These new models enable the identification of disease modifiers that drive specific phenotypes, opening the door for future characterization of protective targets in neural populations most susceptible to neurodegeneration.

## DISCUSSION

SCA17 is caused by abnormal expansion of the polyQ tract in TBP, a highly conserved component of the basal cellular machinery essential for eukaryotic transcription initiation, growth, and development of virtually all cells (Burley and Roeder 1996). It is perhaps not surprising that evidence from animal models suggests that TBP mutation contributes to SCA17 progression in 2 ways: 1) Through toxic gain-of-function, resulting in protein misfolding and neural cell death and 2) Through loss-of-function of endogenous TBP (Nakamura *et al*. 2001; Hsu *et al*. 2014; Toyoshima and Takahashi 2018; Liu *et al*. 2019). We describe here two new *Drosophila* models that ectopically express human TBP harboring polyQ tracts within either the wild-type (Q25) or SCA17 (Q63) disease range, providing novel tools to dissect the relative contribution of these effects to polyQ neurodegeneration.

Human and *Drosophila* TBP share approximately 63% amino acid similarity (Figure 2E), and our data indicate that both our wild-type and polyQ expanded transgenes perturb normal *Drosophila* development. Furthermore, the fact that global developmental (but not adult-specific) TBP expression causes pronounced developmental delay, small pupal size, and severe physiological toxicity in Q25 flies provides a possible explanation for genetic and clinical heterogeneity among SCA17 patients with intermediate CAG/A repeat length. Here, global expression of polyQ-expanded TBP confers high embryonic and pupal mortality, with about 10% of flies eclosing as adults. Interestingly, although a greater number of embryos expressing wild-type TBP survive to larval stages, developmental lethality is more severe, with arrest at the pharate pupal stage and almost no adult flies emerging. It’s formally possible that ectopic wild-type human TBP expression has a dominant negative effect, outcompeting and interfering with endogenous dTbp function essential for normal development. In contrast, the increased aggregation propensity and nuclear accumulation of polyQ expanded TBP may disrupt its normal transcriptional role, resulting in less developmental dysfunction but more severe adult-specific, neurodegenerative effects. Taken together, these findings lend further support to the idea that exogenous TBP overexpression interferes with intrinsic transcriptional control, and that organismal TBP expression is precisely regulated to ensure normal function.

The essential role for TBP in transcription initiation is highlighted by the fact that loss of function in either flies or mice is developmentally lethal (Martianov *et al*. 2002; Hsu *et al*. 2014). Previous investigations further determined that polyQ-expanded TBP binds more tightly with DNA, disrupting intrinsic TBP function and resulting in compensatory loss of wild-type protein (Friedman *et al*. 2008; Huang *et al*. 2015). Our results further support these ideas, underscoring the contribution of dysfunctional TBP to SCA17 neurodegeneration.

We also observe neurodegenerative phenotypes when TBP is expressed specifically in fly eyes, a long-used genetic platform for understanding proteotoxic neurodegenerative diseases (Bonini 1999; Bonini and Fortini 2003; McGurk and Bonini 2012; Tsou *et al*. 2013; Blount *et al*. 2014; Burr *et al*. 2014; Casci and Pandey 2015; Tsou *et al*. 2015; Tsou *et al*. 2016). Here, we notice age- and polyQ repeat length-dependent retinal defects in our Q25 and Q63 flies, allowing for future investigation of genetic or pharmacological modifiers SCA17 neurodegeneration in a robust and easy-to-use genetic model.

Like other polyQ family disease members, neurons are particularly impacted by TBP misfolding, aggregation, and nuclear inclusion despite the mutant disease gene being broadly expressed. By restricting ectopic TBP expression to specific adult tissues, we observed cell-specific contributions to neurotoxicity in our *Drosophila* models. Adult-specific whole-body TBP expression reduced survival independent of repeat length without having significant effects on mobility. On the other hand, Q63 expression in either adult neurons or adult glia was consistently more toxic than Q25 expression in the same tissues, causing reductions in both longevity and mobility. These detrimental phenotypes correlated with higher Q63 aggregation and increased intranuclear inclusions, in agreement with previous findings in animal models and humans (Yang *et al*. 2017; Toyoshima and Takahashi 2018; Liu *et al*. 2019). These data suggest some proteotoxicity from both Q25 and Q63 expression in *Drosophila*, with more pronounced mobility effects when polyQ expanded TBP is expressed in neurons and glia. Neurotoxicity in flies with Q63 expression in adult glia — but not neurons — suggests a cell non-autonomous role of polyQ expanded TBP that warrants further examination.

These new *Drosophila* lines are an important addition to the collective library of genetic tools for polyQ disease models, allowing efficient examination of the biochemical pathways that contribute to SCA17. TBP is universally essential for eukaryotic transcription, and understanding how its function and dysregulation contribute to neurotoxicity in animal models will aid in our understanding of both neural development and pathophysiology. Previous studies have also implicated mutant TBP dysfunction in other polyQ family disorders (Hsu *et al*. 2014), suggesting some common pathways of progressive neurodegeneration in this family of diseases. The curation of these tools, with comparable protein expression levels and identical genetic backgrounds, will allow us to precisely, expediently, and cost-effectively explore both disease-specific and shared pathways driving polyQ neurodegeneration.

## MATERIALS AND METHOD

### Fly Stocks and Maintenance

Gifted stocks used in this study were sqh-Gal4 (Daniel Kiehart, Duke University), elav-Gal4 (GS) (RJ Wessells, Wayne State University), Repo-Gal4 (GS) (Hua Bai, Iowa State Univesity, and w^1118^ (Russ Finley, Wayne State University) and y, w; +; attP2 (Jamie Roebuck, Duke University). To generate transgenic lines that express wild-type (Q25) and polyQ expanded (Q63) TBP through the Gal4-UAS system (Brand and Perrimon 1993), the DNA sequence of full length, human TBP was inserted into pWALIUM10-moe plasmid (DNA Resource Core at Harvard Medical School, MA, USA) with a 3’ HA tag and a construct-specific (25Q or 63Q) polyQ expansion. Constructs were injected by the Duke University Model System Injection Service into y, w; +; attP2. For transgene verification, genomic DNA were extracted from different founder lines using DNAzol (ThermoFisher, Waltham, MA USA) and PCR-amplified using the below primers:

white-end-F: 5′-TTCAATGATATCCAGTGCAGTAAAA-3′
attP2-3L-R: 5′-CTCTTTGCAAGGCATTACATCTG-3′
hsp70-F: 5’-CTGCAACTACTGAAATCTGCCA-3’
pW-ftz-R: 5’-CGTGTGTGATGCCTACCTGA-3’

Prior to all experiments, fly cultures were maintained at a constant density for at least two generations. 20-25 virgin females (depending on genotype) and 5 males were mated in 300 mL bottles with 50 mL standard 10% sucrose 10% yeast spiked with 500 μL Penicillin-Streptomycin (10,000 u/mL, 10mg/mL in 0.9% sterile NaCl (Sigma Aldrich). Adult progeny were synchronized by collecting within 6 hours of eclosion over a 24 hour time period. Groups of 20 age and sex-matched flies were immediately transferred into narrow polypropylene vials containing 5mL of standard 10% sucrose 10% yeast (no antibiotic) or RU486 food as indicated by experiment.

Flies were housed in a 25^°^C incubator on a 12:12h light:dark cycle at 40% relative humidity. Control flies for all non gene-switch Gal4-UAS experiments consisted of heterozygous Gal4 lines backcrossed into y, w; +; attP2 and/or w^1118^ flies, depending on experiment. For gene-switch experiments, RU-flies of the same genotype served as the negative control. RU+ group received 100 μM RU486/mifepristone (Cayman Chemical, Ann Arbor, MI), which activates the gene switch (GS) driver, while RU-group received the same volume of vehicle solution (70% ethanol). Adult progeny were synchronized by collecting within 12 hours of eclosion over a 24 hour time period. Groups of 20 age- and sex-matched flies were immediately transferred into narrow polypropylene vials containing 5mL of standard 2% agar, 10% sucrose, 10% yeast with appropriate preservatives. Food vials were changed every second to third day.

### Developmental biology

Developmental survival was performed by placing 24 gravid female control, Q25 or Q63 flies in aerated 6 oz bottles capped with grape juice agar plates seeded with yeast for 72 hours, after which adults were removed and embryo counted. Embryo were transferred to 6 oz. bottles containing standard 10% sucrose/10% yeast media and allowed to develop at 25°C. Viable adults or arrested pharate pupae were phenotypically scored and counted twice per day.

A similar method was used to score pupariation rate, with the day of embryo transfer considered hour 48 (+/-24h after egg laying, AEL). Twice daily, bottles from each genotype were scored for pupariation until no more larvae emerged. Following pupariation rate assay, a representative sample of at least 10 pharates per genotype were removed and affixed to a glass slide for imaging and measurement at 2X magnification. All pupae were isolated and photographed on the same day. Images were quantified using Image J. 3 individual bottle crosses were performed for each experiment. Statistical analyses were performed in GraphPad Prism (San Diego, CA, USA), specified in figure legends.

### Longevity

At least 400 adults were age-matched and separated by sex within 12 hours of eclosion. Flies were transferred to narrow polypropylene vials containing 5mL of standard 2% agar, 10% sucrose, 10% yeast food. Flies were transferred and scored for death events every 2-3 days until no flies remained. Survival curves were analyzed by log-rank in GraphPad Prism (San Diego, CA, USA). Longevity experiments were performed in duplicate and in parallel with background controls, with each individual graph depicting a representative biological repetition.

### Climbing Speed

Mobility was assessed using the Rapid Negative Geotaxis (RING) assays in groups of at least 100 flies as described in (Sujkowski *et al*. 2022). Briefly, 5 vials of 20 age- and sex-matched flies were briskly tapped down, then measured for climbing distance 2s after inducing the negative geotaxis instinct. For each group of vials, an average of 5 consecutive trials was calculated and batch processed using ImageJ (Bethesda, MD). Flies were longitudinally tested twice per week for 5 weeks or until fewer than 5 flies remained per vial. Between assessments, flies were returned to food vials and housed normally as described above. Negative geotaxis results were analyzed using linear regression in GraphPad Prism (San Diego, CA, USA). Mobility experiments were performed in duplicate, with one complete trial shown in each graph with the following exception:

For both longevity and motility measurements in Figure 3, viability of sqh-Gal4 > UAS TBPQ25 was too low for either parametric assessment of survival or regression analysis of mobility. Mixed-effects models were used to estimate significance, and remaining experiments utilized adult-specific expression to circumvent developmental lethality and obtain necessary statistical power.

### Eye Scoring

Eye scores were represented using the below scoring system, with higher numbers indicating worsening phenotypes: 1) Normal eye with a clear pseudopupil; 2) Diffuse pseudopupil; 3) No visible pseudopupil; 4) Pseudopupil loss accompanied by color variegation/depigmentation at the edge of the eye; 5) Pseudopupil loss accompanied by depigmentation throughout the eye.

### Western Blots

Three whole flies per biological replicate were homogenized in boiling lysis buffer (50 mM Tris pH 6.8, 2% SDS, 10% glycerol, 100 mM dithiothreitol), sonicated, boiled for 10 min, and centrifuged at 13,300xg at room temperature for 10 min. Primary antibody used was rabbit polyclonal Anti-TBP (1:1000, TFIID, Santa Cruz); Secondary antibody: peroxidase conjugated anti-Rabbit (1:5000, Jackson Immunoresearch). Western blots were developed ChemiDoc (Bio-Rad) and quantified with ImageLab (Bio-Rad). For direct blue staining, PVDF membranes were submerged for 10 min in 0.008% Direct Blue 71 (Sigma-Aldrich) in 40% ethanol and 10% acetic acid, rinsed in 40% ethanol/10% acetic acid, air dried, and imaged. Western blots were performed using 5 biological replicates per genotype/sex, and statistical analysis was performed in GraphPad Prism.

### NMJ Histology

NMJ dissections and staining were modified from (Sidisky and Babcock 2020). Briefly, whole flies were anesthetized fly nap then submerged for 30 seconds in 70% ethanol to remove wax coating from cuticle before being transferred to a Sylgard coated dissecting dish. Thoraxes were isolated and fixed in 4% paraformaldehyde for 60 minutes. Following fixation, thoraxes were submerged in liquid nitrogen for 10 seconds, bisected with a sharp razor blade under a dissecting scope, and transferred to ice cold PBS. Samples were then blocked for two hours before staining overnight with primary antibodies (rabbit TFIID, Santa Cruz, 1:100). The following day, tissues were washed, and stained with fluorescent conjugated primary and secondary antibodies (AlexaFluor (AF)640 HRP, 1:200, AF594 phalloidin, 1:1000, anti-rabbit AF488, 1:200, DAPI, 1:10,000ThermoFisher), and imaged using the WSU Department of Physiology Confocal Microscopy Core using a Leica DMI 6000 outfitted with a Photometrics Prime 95B CMOS camera and X-light spinning disc Confocal. Colocalization was quantified using ImageJ using 10 biological replicates per genotype, and statistical analysis was performed in GraphPad Prism.

## Data availability statement

Lines, plasmids and source data are available upon request. The authors affirm that all data necessary for confirming the conclusions of the article are present within the article, figures, and tables.

---

**Supplemental Figure S1:**
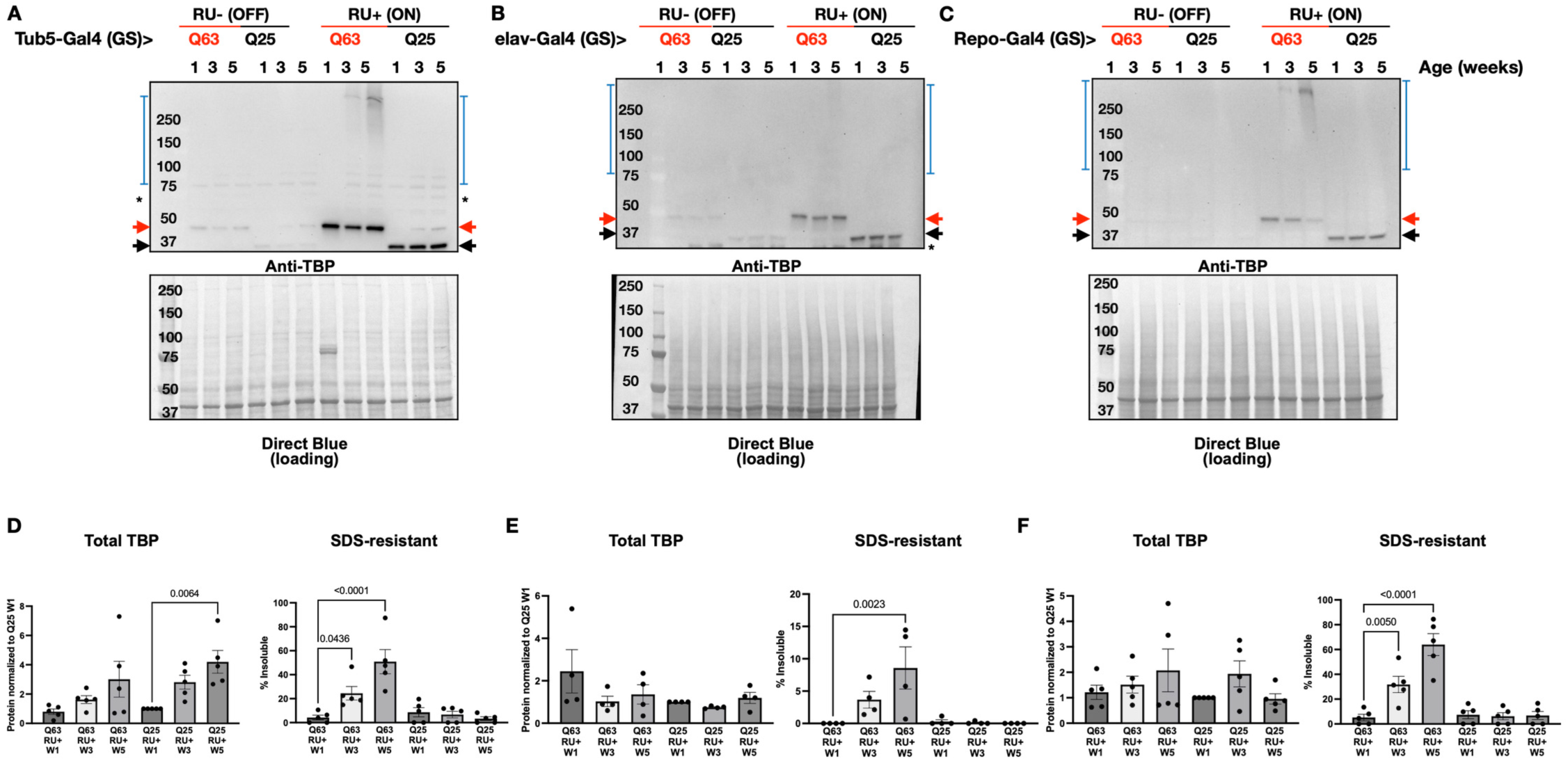
Impact of aging on TBP aggregation in male flies. Representative Western blots in uninduced male control (RU-(OFF)) and experimental flies (RU+ (ON)) with adult-specific pathogenic (Q63, red) or wild-type (Q25, black) TBP expression in weeks 1, 3, and 5 of adulthood, indicated by lane. Expression was induced on adult day 3 in **(A)** all tissues **(quantified in D)**, neurons **(B, E)**, or glia **(C, F)** and continued until sample isolation. Black arrows: TBP, Red arrows: polyQ expanded TBP, Blue brackets: SDS-resistant TBP, asterisks: non-specific signal. Quantification and statistics: ANOVA with Dunnett’s post-hoc comparison. Bars indicate mean -/+ SD, n=5 biological replicates of 3 flies per lysate.

